# Understanding highland adaptation of *Apis cerana* through repeated while independent colonization

**DOI:** 10.64898/2026.02.05.704106

**Authors:** Fushi Ke

## Abstract

Understanding the genetic basis of polygenic adaptation is challenging. Populations that undergo parallel evolution serve as experimental replications enabling the study of molecular mechanism. *Apis cerana* is widely-distributed in Asia and has repeatedly colonized the Qinghai-Tibet Plateau. The highland populations are derived from lowland colonies while showed not admixture with each other, representing independent events of colonization. We investigated resequencing genomes of five populations to study the genetic basis of highland adaptation in *A. cerana*. Using two complementary methods, we isolated genes with adaptive signals in each lineage. Although large proportion of adaptive loci are unique to each population, most potentially adaptive polymorphism shared with genetic variation in lowland populations, consistent with widespread signals of soft selection sweeps. Whereas parallelism was low at the level of adaptive loci, it was greater for functional pathways and greatest for phenotypes. Further enrichment analysis found the shared adaptive loci were overrepresented in pathways related to the development of sensory system and body morphogenesis. These highly connected pathways and loci could buffer different genetic paths and maintain developmental stability. Adaptive signals in the development-related loci suggest stabilizing selection further drive phenotypic convergence under similar stress. However, lineage-specific loci and pathways facilitated adaptive divergence in each lineage within the broadly similar highland environment. Our results demonstrate the genetic redundancy of highland adaptation and lineage-specific evolution via independent colonization in *A. cerana*. This underscores that predicting a population’s adaptive potential requires understanding its full adaptive architecture within the ecological context—including abiotic and biotic interactions.

## 1. Introduction

Recent advances in sequencing technology with plummeting sequencing costs facilitate much works on genome-wide association studies (GWAS), providing insights that most traits are highly polygenic (Csilléry et al., 2018), with contributions of large number of interactive genes outside the core pathways (‘omnigenic’ model, Boyle et al., 2017). This is in aligned with a common finding in evolutionary researches that populations (both natural and laboratory) often possess more adaptive genetic variants than necessary to reach a fitness peak, pointing to significant genetic redundancy in polygenic adaptation (Barghi et al., 2019; Burny et al., 2021; Yeaman, 2022). Since many alternative genetic pathways can produce similar adaptive outcomes, polygenic adaptation is typically heterogeneous and non-parallel at the molecular level. This inherent complexity means that the genetic basis of polygenic adaptation cannot be fully captured by any one-time study without replicates (Barghi et al., 2019).

Parallel evolution, defined as the independent evolution of similar traits in lineages with similar ancestral phenotypes, provides compelling evidence for the deterministic role of natural selection (Cerca, 2023). The extent and pattern of parallelism are governed by a hierarchy of factors. Fundamentally, natural selection in response to common environmental challenges is the primary driver (Cerca, 2023; Yeaman et al., 2016). The complexity of the trait’s genetic architecture is paramount: traits with a simple genetic basis often show higher genetic parallelism (Pickersgill, 2018), whereas polygenic traits frequently evolve through different genetic combinations due to widespread genetic redundancy (Barghi et al., 2019). In addition, the evolutionary history and genetic divergence of lineages critically constrain the outcomes (Sow et al., 2025; Thorhölludottir et al., 2025). The closely related populations possess shared standing genetic variation from a common ancestor could lead to parallel evolution (Barrett and Schluter, 2008; Conte et al., 2012), while lower parallelism observed for populations with differentiated evolutionary paths (Sharma et al., 2022; Storz et al., 2021) and elevated divergent levels (Thorhölludottir et al., 2025).

Although parallelism underlying polygenic adaptation at the genetic level can be heterogeneous, parallelism that manifest at higher hierarchical levels (i.e., functional or phenotypic) may be strong (Allard et al., 2025; Chang et al., 2025). Genome-wide scans often find limited sharing of specific selected loci across populations, yet higher-order convergence may be observed at the level of functional pathways or gene networks (Shakya et al., 2025; Thorhölludottir et al., 2025). This implies that shared evolutionary pressures can drive adaptation through predictable functional channels, even if the precise molecular basis differs. Phenotypic convergence that represents a visible manifestation of the powerful selection (e.g., stabilizing and/or directional selection) in harsh environments could be even stronger since they are tightly linked to similar ecological or functional roles (e.g., Lipshutz et al., 2025; Sackton and Clark, 2019; Swaminathan et al., 2024; Xu et al., 2017; Zhang et al., 2021), and can arise when independently evolving populations experience similar and stable selective pressures (Colosimo et al, 2005; Lipshutz et al., 2025; Pfenninger et al., 2015; Swaminathan et al., 2024; Tishkoff et al., 2007). Beyond these external evolutionary forces, internal component of “selection” due to functionally interacting loci and pathways could maintain phenotypic stability (Schwenk and Wagner 2001) through developmental constraint and canalization (Arnold, 1992; Hallgrimsson et al., 2019; Losos, 2011). Maintaining phenotypic stability is important for functional intergrity of organism against genetic and environmental perturbations (Arnold 1992; Losos 2011; Schwenk and Wagner 2001; True and Haag 2001). External trait selection (e.g., stabilizing selection) from extreme environments will further restrain development by reducing variation on loci and contribute to phenotypic convergence of the lineages under similar stressors (Burny et al., 2021; de Vladar and Barton, 2014; Rosenblum et al., 2014).

The Qinghai-Tibet Plateau (QTP), usually termed as “the Roof of the World”, is well-known for its harsh habitats, such as lower oxygen and temperature, and high intensity of ultraviolet (UV) radiation. Animals inhabited in this area thus evolved with adaptation strategies tackling with the same environmental stress, including more efficient aerobic capacity (e.g., Pamenter et al., 2020; Storz and Scott 2019; Wu et al., 2020), refined thermoregulation and optimized energy metabolism (e.g., Ding et al., 2018; Ellers and Boggs 2004; Parkash et al., 2008), and enhanced resistance to UV radiation (e.g., Wu et al., 2020; Zhang et al., 2021). The evolutionary convergence is pronounced in functionally conserved pathways, such as hypoxia response network (Pamenter et al., 2020; Zhang et al., 2026), even between *Drosophila* and Humans (Babakhanlou 2018; Zhou et al., 2021). However, lineage-specific adaptation strategies evolved in different lineages due to divergent genetic background (e.g., Pamenter et al., 2020; Storz and Scott 2019; Wu et al., 2020; Zhang et al., 2016) and fine-scale environmental pressures (e.g., Chang et al., 2025). For insects, whole body organization, the sum-total of their habits and life-histories constitute their highland specialization (Mani, 1968). Similar external selection would drive phenotypic convergence/ parallelism in different insect lineages/populations, such as the pronounced melanism of highland insects compared with lowland forms (Ellers and Boggs 2004; Mani, 1968; Parkash et al., 2008). Nevertheless, the specialization of highland insects lies in the combination of complex characters that are not only associated with the shared stressors, but also the unique environments not met elsewhere (Chaturvedi et al., 2022; da Silva Ribeiro et al., 2025; Mani, 1968).

The eastern honeybee, *Apis cerana*, is widely distributed in Asia, and has colonized QTP repeatedly from lowland populations (Figure 1A). A recent study (Montero‐Mendieta et al., 2019) based on a pair of highland and lowland populations identified a couple of candidate loci that may mediate high-altitude adaptation of this species. However, these findings should be interpreted with caution. The identified candidates may reflect adaptation to local and site-specific conditions rather than to the core high-altitude stressors. The polygenic nature of highland adaptation further suggests that these genes likely constitute only a subset of the complete adaptive suite. In this study, we used a total of five populations that independently colonized QTP, to investigate the molecular mechanism underlying highland adaptation of *A. cerana*. We tested the hypothesis that these populations share loci and pathways facilitating adaptation to core highland stressors, while also possess lineage-specific loci and pathways adapted to the local environment.

**Figure 1.**
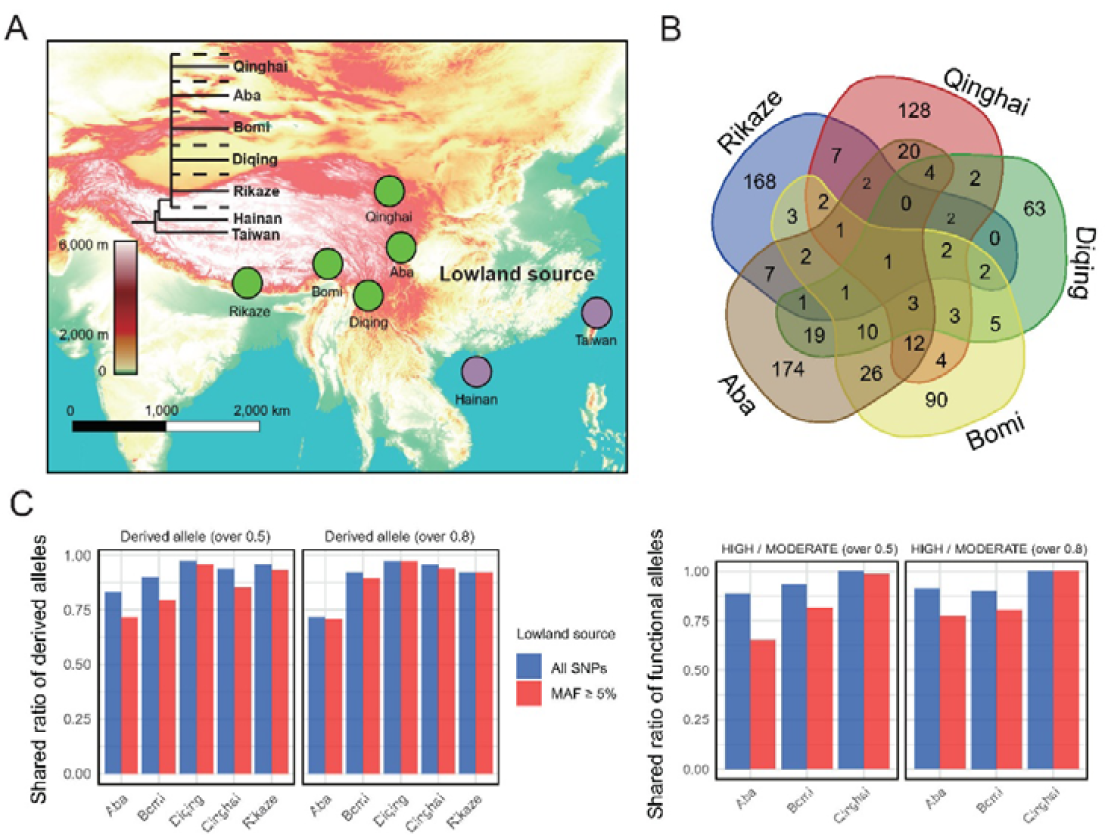
Characterization of adaptive polymorphism in *Apis cerana* highland populations. **(A)** Geographic distribution and phylogeny of *A. cerna* populations. Five highland populations are labelled with circles of green color. Two lowland populations (purple circles) are used as control in selection sweep analysis using haplotype homozygosity statistic. All lowland source samples are from Central lineage of *A. cerana* Lineage M (Ke and Vessur, 2026) and collected at locations with elevation lower than 2000 meters. A simplified phylogenetic relationship was inferred from Ke and Vasseur (2026). The dash lines represent populations/individuals from the lowland; **(B)** Venn plot showing shared and unique adaptive genes in each highland population; **(C)** Ratio of potentially adaptive polymorphism in highland populations shared with lowland Central lineage. HIGH and MODERATE represent variants with functional impacts (annotated with SNPeff) on adaptive genes.

## 2. Materials and Methods

### 2.1 Identification of selective signals in *Apis cerana* highland populations

Asian honey bee (*A. cerana*) has repeatedly colonized highlands many times and inhabited at different geographical locations with high elevation (Figure 1A, with elevation higher than 2000 m). At least five different populations, including Aba, Bomi, Diqing, Qinghai, and Rikaze, have successfully colonized Qinghai-Tibet plateau (Figure 1A and Supplementary Table 1). They are all derived from the central lineage of *A. cerana* Lineage M recently (less than 700 thousand years, Ke and Vassuer, 2026) while show no admixture with each other (Figure 1A, Ji et al., 2020; Ke and Vassuer, 2026). There are two conflicting models explain polygenic adaptation. Population genetics views it as a series of selective sweeps at single loci, whereas quantitative genetics sees it as a collective response achieved through allele frequency shifts across many loci (e.g., Höllinger et al., 2019). Nevertheless, recent theoretical and empirical studies have shown both models could result in a sweep-like genome signature (Barghi and Schlötterer, 2020), with detectable signals in local haplotype blocks (Höllinger et al., 2019). With the aim to characterize genetic variation underlying adaptation of *A. cerana* populations to highland habitats, we employed the haplotype homozygosity statistic H12 that could detect hard as well as soft sweeps (Garud et al., 2015) to identify adaptive signals. In addition, we used the site frequency spectrum (SFS) method in SweepFinder2 (DeGiorgio et al., 2016) in adaptive gene isolation as a complementary analysis.

Individuals from each population/lineage were isolated from the *A. cerana* resequencing dataset (Ke and Vassuer, 2026) with Vcftools (v 0.1.16, Danecek et al. 2011) and phased using beagle (beagle.28Jun21.220.jar, Browning et al., 2021) with default parameters. For each population, the input data of each chromosome for H12 analysis was prepared with Formtools command in iTools (github.com/BGI-shenzhen/Reseqtools/), and ran with H12_H2H1.py script (github.com/ngarud/SelectionHapStats/) in a window size of 400 SNPs and 25 SNP step following the manual, which use *Drosophila melanogaster* data as an example. Since *A. cerana* has a recombination rate near 20 cM/Mb (Shi et al., 2013), which is much higher than the *D. melanogaster* (< 5 cM/Mb, Wang et al., 2023), we further tested smaller window sizes (100 and 200 SNP window) to investigate the selection sweep signals. Window with the top 1% of the genome-wide H12 values were isolated. We further identified genes that are overlapped with the regions with selection sweep signal (i.e., overlapped with the 10-kb window of the selection signal).

For SweepFinder2 analysis, one individual from *A. cerana* Lineage S (Su et al., 2023) was used as the outgroup to polarize each site. The folded Site Frequency Spectrum (SFS) was then generated with --counts2 in vcftools (v 0.1.16, Danecek et al. 2011). With genome-wide SFS as the reference, we scanned empirical SFS in each window and isolated selection sweep signals with a user-defined number of 1000 grid points in each chromosome. We then identified closely related genes as adaptive candidates (genes overlapped with the 10-kb window of the selection signal). It remains difficult to identify and verify the causal locus (e.g., SNP) of the gene underlying a monogenic or polygenic trait, which needs in depth molecular validation or quantitative trait locus (QTL) mapping.

Therefore, instead of pinpointing to causal region of the adaptive gene (e.g., untranslated or coding region) or identifying the causal variant (e.g., SNP), our strategy here is to isolate genes closely related to the adaptation (i.e., with sweep signals) and provide a more comprehensive while robust (with replications) adaptive gene set underlying the complex traits.

### 2.2 Filtering adaptive genes in each highland population

To have a list of adaptive genes with high confidence, we used two methods in isolating genes with selection signals. First, overlapped genes (named as A1) in SweepFinder2 gene set (related to regions with top 1% CLR) and the list identified based haplotype homozygosity statistic H12 (either from 100, 200 or 400 SNP window size) were deemed as adaptive genes. Second, overlapped genes that were identified only based on H12 statistic. Since our earlier *A. cerana* study (Ke and Vasseur, 2026) confirmed that reduced genetic shuffling from genome features like low recombination would let some genomic regions more prone to accumulating linkage-based sweep signals, such as H12. We thus employed two populations (Hainan and Taiwan populations of *A. cerana* Lineage M, with 25 and 12 individuals respectively, Supplementary Table 1) from lowlands and with different level of nucleotide diversity (Ke and Vasseur, 2026) as control to remove potentially pseudo selection sweep signatures identified in haplotype homozygosity statistic. We first investigated the adaptive loci presented in at least two of the three H12 analyses (i.e., 100, 200, 400 SNP window size, named as B0) of the highland populations and two lowland controls. Further, genes (in B0) identified based on H12 in each highland population that were overlapped with either of two lowland populations (B0) were deemed to be noisy signals and excluded for further analysis of highland adaptation. The filtering will remove some true signals but would increase the confidence of adaptive targets. With this filtering, we have an adaptive gene list (B1) for each highland population solely based on H12 statistic. By combining SFS parameter (SweepFinder2 analysis) and haplotype homozygosity statistic (H12), we have an adaptive gene list (merging A1 and B1) for each highland population (filtering pipeline in Supplementary Table 2).

### 2.3 Enrichment analysis for the adaptive loci

Three datasets were used for further enrichment analysis: 1) gene list in each of five populations; 2) shared gene list in at least two populations; 3) all gene pool in five populations. We characterized the adapted genes with Gene Ontology (GO) analysis in two ways. We first performed GO analysis in ClusterProfiler (v 4.0, Wu et al., 2021) with the *Drosophila melanogaster* database (BDGP6.46) as reference (i.e., org.Dm.eg.db). *Drosophila melanogaster* is the model species with the best annotation in insects, and thus will provide the most robust evidence for insects in conserved pathways, such as GO terms related to development. Second, we conducted enrichment analysis with the *A. cerana* gene annotation database as a complementary. We annotated all *A*.*cerana* gene list (AcerK_1.0, GCF_029169275.1) by blasting against the eggNOG 5.0 database (Huerta-Cepas et al., 2019) using eggNOG-mapper (v2.1.12) (Huerta-Cepas et al., 2017) and conducted GO analysis with clusterProfiler (v 4.0, Wu et al., 2021) to identify the significantly (p.adj <0.05) enriched GO terms. In this analysis, non-insect GO terms were removed manually. R package ggplot2 (v3.3.3) (Wickham, 2016) was used in generating bubble plot of the significantly enriched GO terms. To visualize functional enrichment results, we constructed and summarized the linkage of enriched concepts (i.e., emapplot) with clusterProfiler. Default settings were used in the mentioned R packages and software.

### 2.4 Characterization the origin of adaptive polymorphisms

Genetic variation underlying adaptation could be resultant from *de novo* mutations (Bolnick et al., 2018) or collateral evolution due to standing genetic variation or introgression (Cresko et al. 2004; Reed et al., 2011; Stern, 2013). To further understand the source of adaptive polymorphism underlying highland adaptation, we calculated percentage of the shared polymorphism between highland and lowland source population (173 individuals, Supplementary Table 1). First, we investigated loci with high derived allele frequency (≥0.5 and ≥0.8) in each highland population. Second, we checked loci with high frequency (≥0.5 and ≥0.8) of functional alleles (with MODERAT and HIGH impacts annotated with SNPeff, Cingolani et al., 2012). In these two steps, we calculated percentage of overlapped polymorphism in each highland population with lowland source (all SNPs and loci with minor allele frequency (MAF) ≥ 0.05). Since all five populations independently colonized Qinghai-Tibet Plateau and are not admixtured with each other, we focused on two mechanisms, that are *de novo* mutations and standing genetic variation, in explaining the pattern.

### 2.5 Phenotypic data analysis

#### 2.5.1 Phenotypic data of the study populations

From Zhang et al. (2025), we obtained morphological measurements from five populations at two elevational gradients: four high-altitude populations (Maerkang: Aba; Linzhibomi: Bomi; Diqingweixi: Diqing; Xunhua: Qinghai) and one low-altitude reference population (Shandong). A total of 40 morphological traits were measured, including linear measurements of skeletal elements (e.g., femur length, tibia length), wing dimensions, and other morphological features. Data from Zhang et al. (2025) were formatted as “mean±SDletter” with letters indicating significance in Tukey HSD tests (Bonferroni method was used for correcting the p-value in multiple comparisons). The following analyses of phenotype were performed in R (version 4.3.0+, R core team, 2023) using tidyverse (Wickham et al., 2019) and ggplot2 (v3.3.3) (Wickham, 2016).

#### 2.5.2 Coding scheme for parallel evolution analysis in phenotype

We recoded each trait for parallel evolution analysis. A three-stage coding procedure was implemented to identify parallel evolutionary patterns:

##### Stage 1: Mean-based coding with significance threshold

Each high-altitude population was compared to the low-altitude control for each trait:

- Code 1: Mean value significantly higher than control
- Code -1: Mean value significantly lower than control
- Code 0: No significant difference

Traits in two populations (i.e., highland population v.s. lowland population) were considered significantly different if they shared no common letters in their annotations (Zhang et al., 2025).

##### Stage 2: Negative code conversion rule

To account for directional consistency in evolutionary change and compare with lineage-specific genes (in 0,1 coding) and GO concepts (in 0,1 coding), we further conversed trait with negative code value:

- If a trait exhibited three or four code -1 values across the four high-altitude

populations, all -1 codes were converted to 1, and the rest population (three populations with -1) will be 0 not matter the value in Stage 1

- We don’t have traits encoding two -1 and two 1 in four highland populations
- This transformation recognizes consistent reduction in trait values across multiple populations represents a parallel evolutionary pattern

##### Stage 3: Phenotypic stability adjustment

To capture parallel evolution in phenotypic stability:

- For each population-trait combination, if the standard deviation (SD) was less than 50% of the control population’s SD, the code was forced to 1
- This adjustment identifies traits where high-altitude populations show reduced phenotypic variation (increased developmental stability), regardless of mean differences

The final binary matrix represented each trait’s evolutionary pattern across the four high-altitude populations relative to the low-altitude control. We filtered the traits with 0 in four populations for further analysis (seven traits were removed). The trait coding matrix was in binary format where:

- Code 1 = Presence of evolutionary change (either increased mean, decreased mean with consistency rule, or reduced variation)
- Code 0 = Absence of evolutionary change or not in the same direction with the other three highland populations

### 2.6 Jaccard Similarity Analysis

We then used adaptive gene matrix, GO concept matrix and trait matrix to investigate parallel evolution at different levels. Pairwise Jaccard similarity indices were calculated between all high-altitude population pairs using the formula:

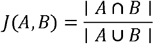

Where:

- *A* and *B* are sets of genes/GOs/traits coded as 1 for populations A and B, respectively
- | *A* ∩ *B* | = Number of genes/GOs/traits with code 1 in both populations
- | *A* ∪ *B* | = Number of genes/GOs/traits with code 1 in either population

To distinguish genuine evolutionary patterns from random associations, we conducted Monte Carlo simulations (10,000 iterations) for each analysis level. Random null distributions were generated by sampling from the combined gene/GO/trait pool while preserving each population’s observed set size. Statistical significance was evaluated through empirical p-values, with convergence and divergence tested separately by comparing observed Jaccard indices to their respective null distributions using a significance threshold of p < 0.05.

### 2.7 Genetic network analysis

Based on the gene sets associated with 13 body development-related Gene Ontology (GO) terms (Figure 2C, obtained from AmiGO2 for *Drosophila melanogaster*, https://amigo.geneontology.org/amigo), we constructed an integrated genetic interaction network. First, each GO-specific gene list was submitted to the STRING database (v11.5, https://cn.string-db.org/) to retrieve protein–protein interaction networks (minimum required interaction score was set as default: 0.4). All GO networks were then merged, retaining only the interaction with the highest combined confidence (i.e., combined_score) for each unique gene pair to remove redundancy. The unified network was further visualized and analyzed using Cytoscape (v3.10.4, Shannon et al., 2003). Nodes with colored pie chart in the network were further used to indicate whether a gene was identified as adaptive in each of the five surveyed populations (Aba, Bomi, Diqing, Original, and Rikaze). We also calculated Degree of each gene (node) in this network. Finally, to examine the core adaptive architecture, we extracted only those genes under selection in each population from the thirteen GO sets and reconstructed a subnetwork representing shared functional connections among adaptive loci. This approach allowed us to assess how developmental network structure enables multiple genetic routes toward phenotypic convergence.

**Figure 2.**
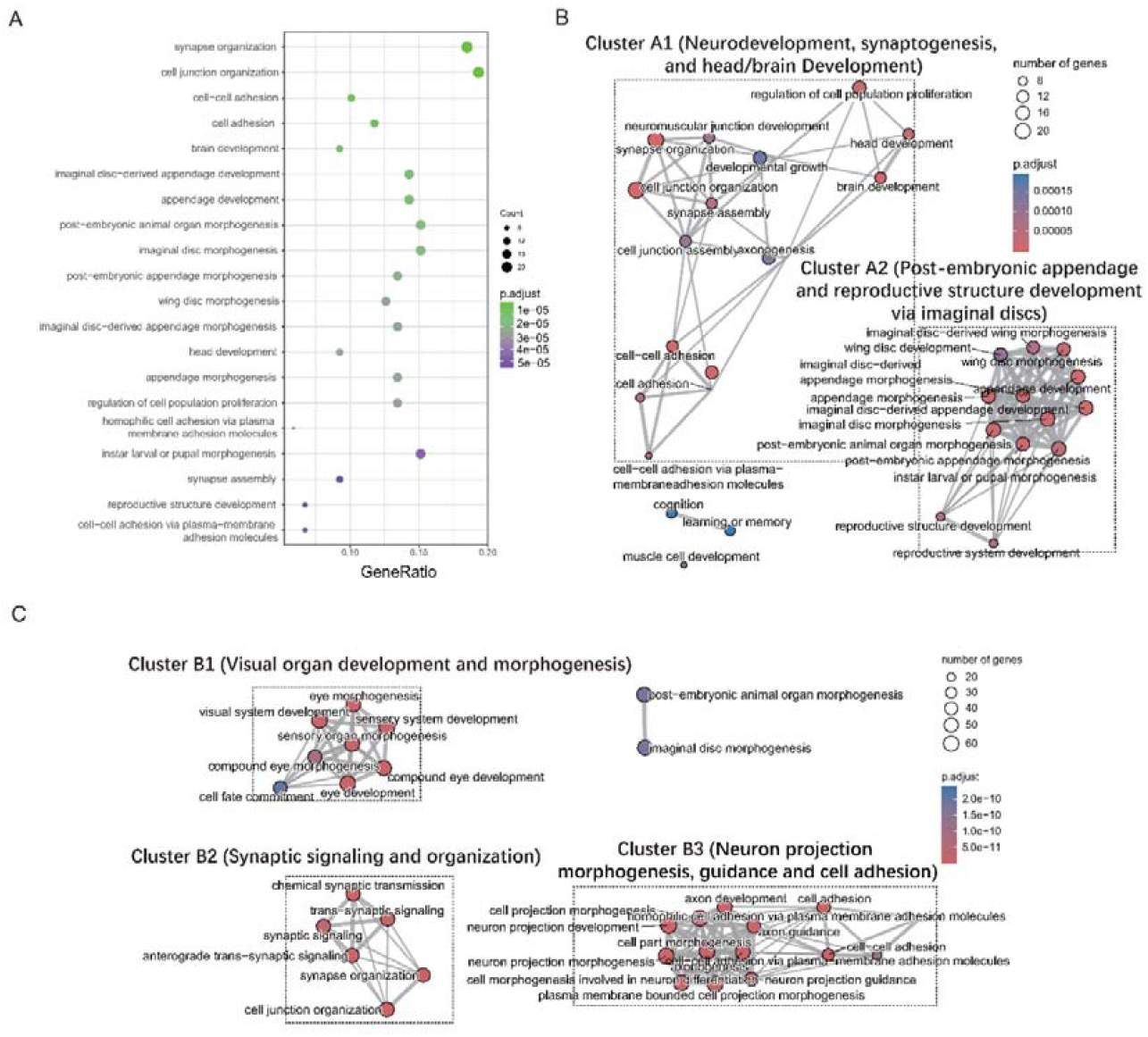
Enrichment analysis of the adaptive loci. **(A)** Top 20 significantly enriched Gene ontology (GO) terms of the adaptive genes (shared in at least two highland populations). Full lists of the significantly enriched GO terms of shared and full adaptive gene list refers to Supplementary table 2 and 3; **(B)** Summary analysis with emapplot shows the relationship of significantly enriched GO terms of shared gene set. Two clusters (Cluster A1, neurodevelopment, synaptogenesis, and head/brain development; Cluster A2, post-embryonic appendage and reproductive structure development via imaginal discs) were in rectangles with dotted line; **(C)** emapplot shows three clusters of the significantly enriched GO terms with all adaptive loci. Three clusters were in rectangles with dotted line.

## 3. Results

### 3.1 Adaptive loci in *Apis cerana* highland populations

We investigated five highland populations (altitude > 2000m) that independently colonized Qinghai-Tibet Plateau for understanding high-altitude adaptation of *A. cerana* (Figure 1A; Supplementary Table 1). Two lowland populations (altitude < 2000m, Figure 1A; Supplementary Table 1) from Hainan and Taiwan Islands were used as controls in removing potentially “adaptive” loci that may be a result of genomic architectures (see Methods). In addition, a total of 178 individuals collected from habitats lower than 2,000 meters across the mainland (Supplementary Table 1) were used as the source population (of the highland population) to investigate possible origin of the adaptive variants. Based on two complementary methods, we identified 283, 167, 118, 193 and 201 genes with adaptive signal in Aba, Bomi, Diqing, Qinghai, Rikaze (Supplementary Table 2 and 3). Most of the adaptive genes are lineage-specific (Aba: 174/283, Bomi: 90/167, Diqing: 63/118, Qinghai: 128/193, 168/201, Figure 1B) with few of these loci shared among populations (e.g., only one gene, *sli*, shared in five populations). We further calculated shared proportion of derived alleles (with derived allele frequency ≥0.5 or ≥0.8) in each highland population with lowland source, and found most of the highland populations exhibited a high degree (over 75%) of common derived alleles with lowland source, with only Aba population had a ratio slightly lower than 75% (Figure 1C). We further investigated derived alleles (with derived allele frequency ≥0.5 or ≥0.8) with potential functions (i.e., HIGH / MODERATE in SNPeff annotation). Most of the populations displayed a high degree of common functional elements with lowland source (higher than 75%), with only Aba has a moderate ratio of 64% (Figure 1C). The high level of allele sharing between highland population and lowland source population support an evolutionary trajectory of adaptation likely from standing genetic variation, a pattern that was consistent with widespread adaptation through soft selection sweeps (Supplementary Table 2).

### 3.2 Enriched pathways facilitate *A. cerana* highland adaptation

For adaptive genes presented in at least two populations, we identified 231 GO terms with an adjusted p valule (p.adj) <0.05 (Supplementary Table 4) (mapping to *Drosophila melanogaster* database unless otherwise stated) in enrichment analysis. Investigation of the first 20 GO terms shows most of the pathways are related to neural development (e.g., synapse organization), and body morphogenesis (e.g., appendage development) (Figure 2A, Supplementary Table 4). Summary analysis of all significantly enriched GO terms with emapplot supports the importance of pathways related to neurodevelopment, synaptogenesis, and head/brain development (Cluster A1), and post-embryonic appendage and reproductive structure development via imaginal discs (Cluster A2) (Figure 2B). These two clusters are with 14 (Cluster A1) and 13 (Cluster A2) highly connected GO terms, respectively (Figure 2B). With all adaptive genes, we identified a total of 539 GO terms (Supplementary Table 5). Summary analysis of these GO concepts with emapplot shows three clusters of GO terms (Figure 2C), including Cluster B1 (visual organ development and morphogenesis, 8 GO terms), Cluster B2 (synaptic signaling and organization, 6 GO terms), and Cluster B3 (neuron projection morphogenesis, guidance and cell adhesion, 14 GO terms). We further conducted GO analysis for each population and investigated shared and lineage-specific GO terms. A total of 30 shared GO terms were identified across five populations (Supplementary Table 6), of which 11 concepts were related to sensory system development (5 concepts are related to eye development) and 10 of them were associated with the morphogenesis of different body parts (Supplementary Table 6). The list of shared GO terms could be found in significantly GO list of enrichment analysis either with shared genes (Supplementary Table 4) or with all adaptive genes (Supplementary Table 5), suggesting the importance of development of sensory system and body morphogenesis during *A. cerana* adaptation in highland habitats.

### 3.3 Convergence of morphological phenotype through directional and stabilizing evolution

We further investigated parallel evolution of five highland populations at gene, GO term and morphological phenotype levels. Although the employed traits of the phenotype (i.e., 40 morphological traits) should have underestimated the convergence (e.g., without neuronal and physiological traits), the Jaccard Index (JI) between each population showed a gradually increased pattern from gene (mean JI: 0.076), GO pathways (mean JI: 0.25) to phenotype (mean JI: 0.39) (Figure 3A). Investigation of the 33 traits (seven traits were removed due to not significant difference in any of highland populations with lowland population) in four highland populations showed 18 of them had mean values all higher or lower than the lowland reference (i.e., Shandong) (Supplementary Table 7), with three traits all significantly differentiated with Shandong population (pigmentation of tergite 2 (PT2), pigmentation of labrum 1 (PLAB1), and pigmentation of labrum 2 (PLAB2), Supplementary Figure 1). This is consistent with phenotype convergence through directional selection. In addition, three traits in all four highland populations showed significantly reduced variation (i.e., > 50% variation) compared with Shandong population (tergite 4, longitudinal (T4), sternite 6, longitudinal (S6L), sternite 6, transversal (S6T), Figure 3B), indicating parallel evolution through stabilizing selection in these highland populations. Although we lack morphological data for Rikaze population, our analysis of the morphometric data as reported by Zhang et al. (2025) reveals a broader pattern: highland populations (encompassing 16 sites on the Qinghai–Tibetan Plateau) show significantly reduced variation in T4, S6L, and S6T compared to lowland group.

**Figure 3.**
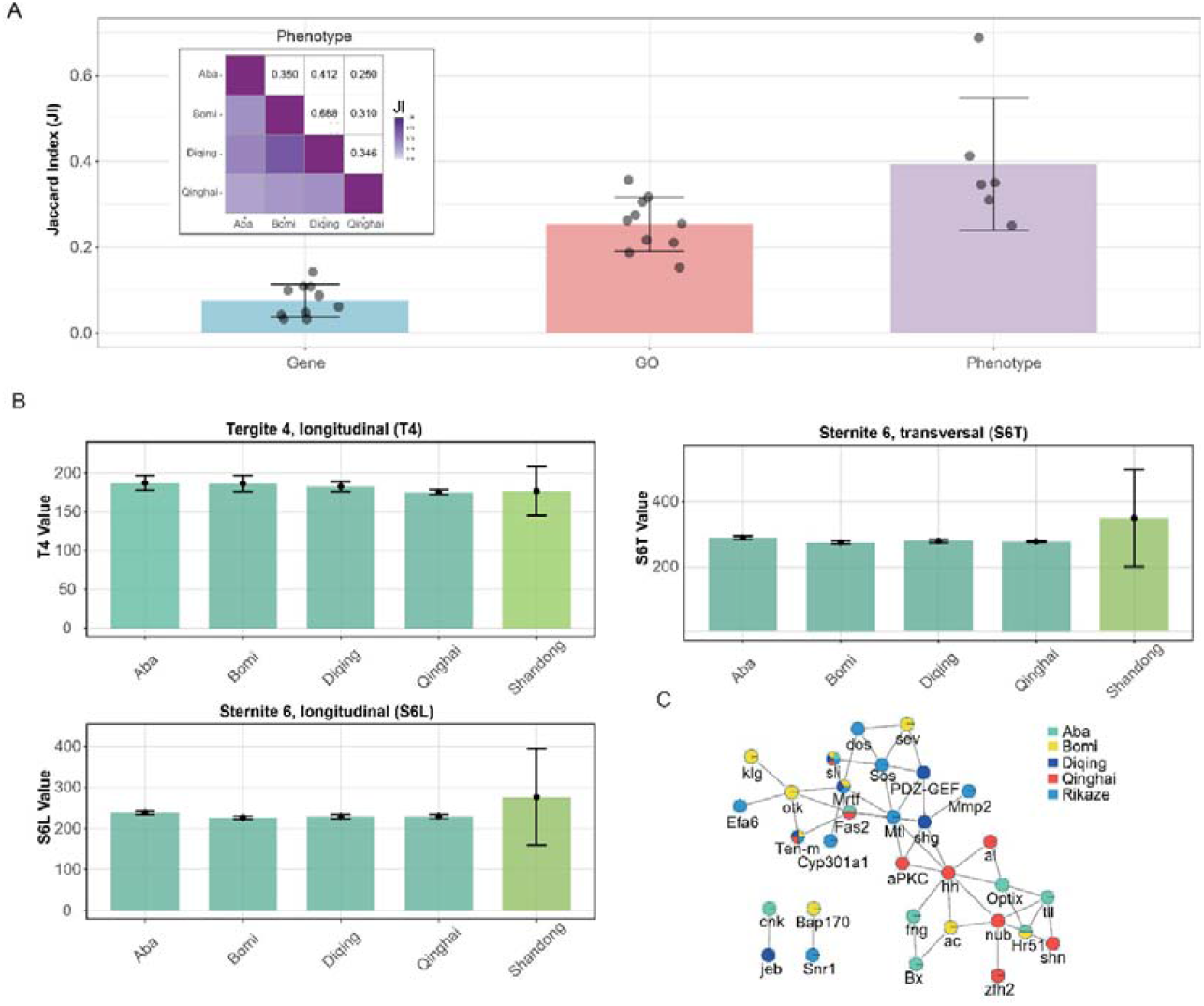
Network buffering enables multiple genetic paths to developmental stability. **(A)** Similarity of five highland populations at genetic, functional, and phenotypic levels. Heatmap depicts phenotypic similarity (Jaccard Index, JI) among four populations (Aba, Bomi, Diqing, Qinghai). Bars represent similarity at the levels of adaptive genes, Gene Ontology (GO) terms, and phenotypes. Higher JI values indicate greater parallelism; **(B)** Phenotypic convergence through developmental stability. Relative uniformity within highland populations compared with lowland reference suggests stronger stabilizing selection toward similar phenotypic optima. The measured value is 100 times of the actual value (mm) (Zhang et al., 2025); **(C)** Adaptive genetic network underpinning developmental stability across populations. Although adaptive genes vary among populations, each population targets multiple genes within this conserved development network. The adaptive loci, *sli*, shared across five highland populations.

### 3.4 Genetic interaction maintains developmental stability

We then investigated adaptive loci related to body development (Cluster A2), one of the two GO clusters previously identified (Figure 2B). All genes in a total of 13 GO terms were downloaded from AmiGO2, and investigated with STRING and Cytoscape. Genes (nodes) in this genetic network (cluster) are highly connected (Supplementary Figure 2), with an average degree (number of direct connections that a gene has with other genes in the network) of 19.46 (Supplementary Figure 3). For adaptive loci, a subnetwork with 27 genes were identified (Figure 3C). Each highland population had several adaptive genes (shared or unique) presented in this network (Figure 3C). Monte Carlo simulations (Supplementary Figure 4) show random accumulation of these adaptive loci in each population (loci in Rikaze likely interacted with other pathways that mediated local adaptation, see below). Although different set of adaptive loci related to body development in each population, the highly inter-connected genetic network could buffer different genetic paths and maintains developmental stability of different highland *A. cerana* populations. Only one adaptive locus, *sli*, was shared across all five populations (Figure 3C) and showed high degree in the developmental network, suggesting its important role as a hub gene in maintaining developmental stability.

### 3.5 Non-random accumulation of adaptive loci and pathways suggest different adaptation strategy in each highland population

We further conducted Monte Carlo simulations at gene, GO terms and phenotype levels to test if the convergence appears with random or not. Null distributions were created by randomly sampling the combined data while keeping the size of each original set constant. Unlike GO terms and phenotype traits (Supplementary Figure 5), simulation of the adaptive gene pool indicated the observed gene set in each population was non-randomly accumulated and differentiated with each other (with JI significantly lower than expected, Figure 4A). This is consistent with differentiated habitats (Figure 4B) and independent colonization (Figure 1A) that leads to accumulation of distinct adaptive loci.

**Figure 4.**
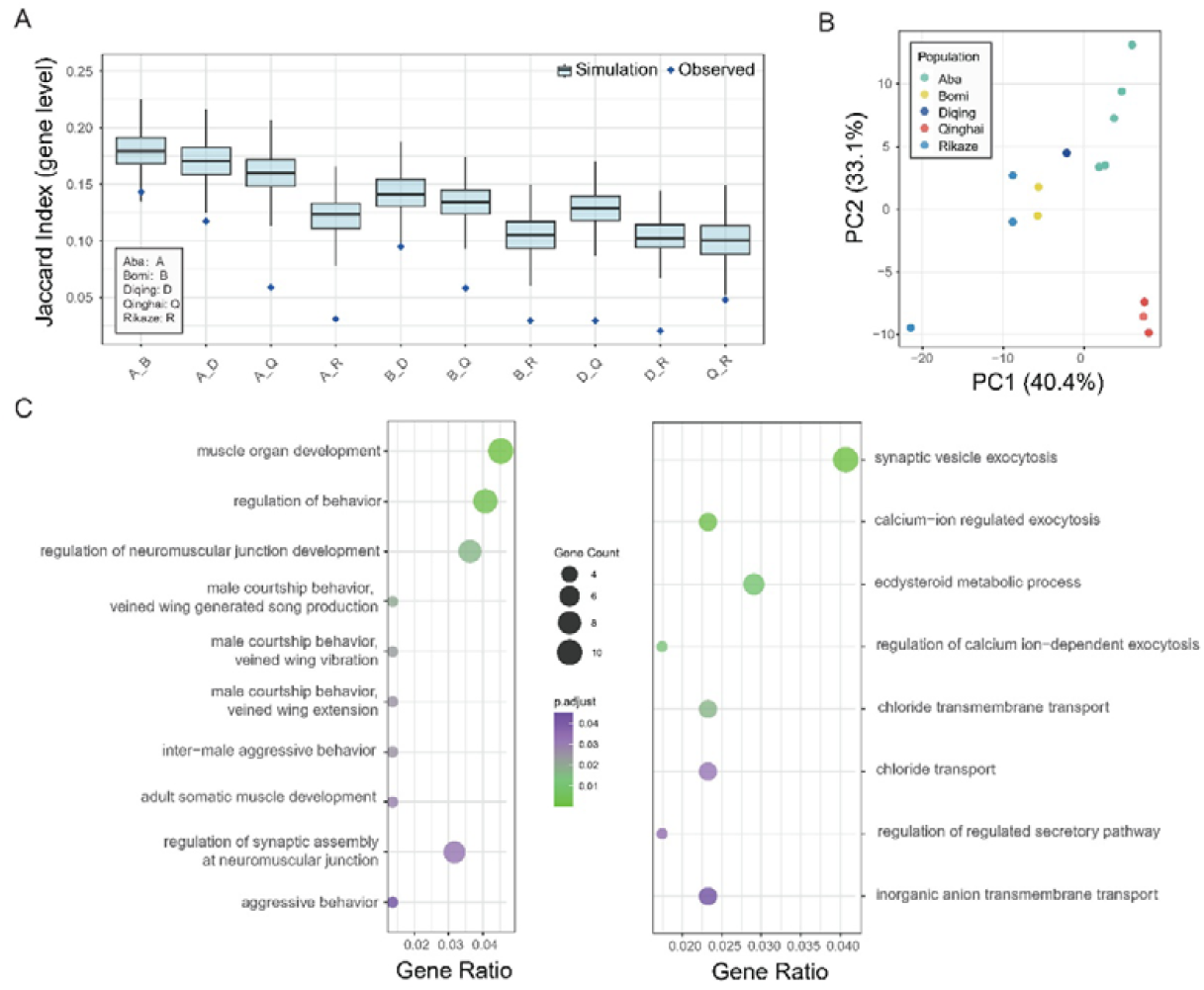
Lineage-specific loci and pathways suggest different adaptation strategies across highland populations. **(A)** Adaptive gene sets exhibit significantly reduced similarity across population pairs. Jaccard Index at the gene level was calculated across five populations (Aba, Bomi, Diqing, Qinghai, Rikaze) and compared to a null distribution generated by 10,000 Monte Carlo simulations. The observed similarity is significantly lower than random expectation, indicating lineage-specific genetic networks and adaptation shape the retention of adaptive alleles across populations; **(B)** Cluster analysis of bioclimatic variables showed environmental differentiation of the highland populations. Full bioclimatic data can be found in Supplementary Table 1; **(C)** Lineage-specific pathways indicate differentiated adaptation strategy. Left panel: selected lineage-specific GO terms in Aba, Right panel: selected unique GO terms in Rikaze.

We next analyzed lineage-specific GO terms of each highland population from differentiated habitats (Figure 4B). A total of 93, 26, 29, 42, and 81 lineage-specific GO terms were identified in the Aba, Bomi, Diqing, Qinghai, and Rikaze populations, respectively (Supplementary Table 6). In the Aba population, apart from muscle organ development, several GO terms associated with male mating behavior were enriched, including GO:0016545 (male courtship behavior, veined wing vibration), GO:0045433 (male courtship behavior, veined wing generated song production), GO:0048065 (male courtship behavior, veined wing extension), GO:0050795 (regulation of behavior). These terms were also supported by GO annotations in the *A. cerana* database (Supplementary Table 8). In Rikaze worker bees, we identified lineage-specific GO terms linked to ion transport and neurosecretory regulation, including GO:0098661 (inorganic anion transmembrane transport), GO:0045455 (ecdysteroid metabolic process), GO:0006821 (chloride transport), GO:0017156 (calcium-ion regulated exocytosis), GO:1903305 (regulation of regulated secretory pathway), GO:0017158 (regulation of calcium ion-dependent exocytosis), GO:0016079 (synaptic vesicle exocytosis), GO:1902476 (chloride transmembrane transport), all of which were supported by annotation with *D. melanogaster* and *A. cerana* databases (Supplementary Tables 6 and 8). In the rest three populations, we didn’t find interesting pathways shared between annotation with *D. melanogaster* and *A. cerana* databases (Supplementary Tables 6 and 8). This is likely due to the divergence of *D. melanogaster* and *A. cerana*, and species-specific evolution in honey bees. Nevertheless, we found lineage-specific GOs terms in each of the highland population not matter which database was used. The non-random accumulation of adaptive alleles and lineage-specific functional pathways across all five populations support that each colony have evolved with distinct adaptations (Figure 1B and Figure 4A) to its fine-scale habitat within the broadly similar highland environment.

## 4. Discussion

Understanding the genetic basis of adaptation is a central goal in evolutionary biology. However, the adaptive variation identified in any case-control comparison represents only a piece in the complex puzzle of polygenic adaptation (Barghi et al., 2019; Yeaman, 2022), underscoring the critical need for replication to unravel its underlying mechanisms. In this study, we investigated the molecular mechanism underlying highland adaptation of *Apis cerana* by analyzing the resequencing genomes of five populations that independently colonized Qinghai-Tibet Plateau (QTP) and showed no mutual admixture (Figure 1A). We verified that under the broadly similar environmental stressors, directional and stabilizing selections have driven phenotypic convergence across highland populations. Shared functional pathways and adaptive loci further highlight the importance of developmental stability (i.e., development of sensory system and body morphogenesis) when confronting with QTP’s harsh conditions. Meanwhile, each highland population possesses lineage-specific loci and pathways, reflecting their unique adaptive strategies to fine-scale environmental heterogeneity. Together, these findings through replicated populations provide a more comprehensive understanding of highland adaptation in *A. cerana*. Finally, the observed genetic redundancy of polygenic adaptation and the prevalence of site-specific adaptation call for cautions in predicting adaptive potential to novel environments and inform strategies for biodiversity conservation in the genomic era.

### 4.1 Highland adaptation of *A. cerana*

Phenotypic convergence (or parallelism) of lineages provides compelling evidence of natural selection (Cerca, 2023). This is because such repeated outcomes are tightly strongly associated with similar environmental stress or functional roles (e.g., Lipshutz et al., 2025; Sackton and Clark, 2019; Swaminathan et al., 2024; Xu et al., 2017; Zhang et al., 2021) and are highly unlikely to arise by chance alone (Cerca, 2023). In *A. cerana*, phenotype convergence of highland population was observed in multiple traits, such as increased pigmentation tergite 2 (PT2), labrum 1 (PLAB1), and labrum 2 (PLAB2). The melanism of body and wing is important for thermoregulation of insects (e.g., Ellers and Boggs, 2004; Parkash et al., 2008). Therefore, we hypothesize that increased pigmentation may have played a key role in the adaptation of *A. cerana* to the QTP environment. In addition, we found parallelism in reduced variation of body development (Figure 3B) across the investigated highland populations when compared with the lowland forms. This supports strong stabilizing selection on development under environmental stress (Burny et al., 2021; de Vladar and Barton, 2014; Rosenblum et al., 2014). A broader investigation of 16 *A. cerana* colonies from the QTP confirms a similar pattern of reduced variation in developmental traits (Zhang et., 2025). This finding demonstrates the importance of developmental stability for highland adaptation in this species. Although our phenotypic data were limited to morphology and lack neuronal or physiological dimensions (some of which may show convergence, see discussion below), the parallel evolution of these morphological traits across populations strongly suggest their adaptation to the similar highland stressors.

Functionally enriched pathways further support parallelism in development across highland populations. GO enrichment analysis revealed a common set of GO terms related to neural development and body morphogenesis across all highland populations. This was further evidenced by the enrichment analysis of the shared gene list, from which we found two GO clusters that are related to development of sensory system and body morphogenesis (Figure 2B). Interestingly, the only adaptive locus that shared across five populations was *sli* that involved in both processes (Brose et al., 1999; Rothberg et al. 1990; Wong et al., 2002; Wu et al., 1999). These development-related loci carried signals of adaptation, supporting strong stabilizing selection in harsh habitats would restrict developmental variation to optimize fitness by reducing genetic variation at these adaptive loci (de Vladar and Barton, 2014; Rosenblum et al., 2014; Lai et al., 2025). Furthermore, these loci are highly interconnected within the gene network (Figure 3C and Supplementary Figure 2) and occupy central positions within it (Supplementary Figure 2), suggesting they are highly pleiotropic. Stabilizing selection and directional selection on such pleiotropic loci thus could lead to parallel evolution responses across populations (Hämälä et al., 2020; Lai et al., 2025).

In contrast to the broadly similar highland stressors, local environments composed of distinct environmental factors could shape a unique suite of adaptive traits (Feng et al., 2024; da Silva Ribeiro et al., 2025). When investigating the adaptive loci, we found lineage-specific enriched GO terms and a nonrandom accumulation of these loci in each highland population. The loci in these pathways may facilitate local adaptation through a combination effect of genetic interaction (epistasis or pleiotropy) and gene-environment interaction. In the Rikaze population, we identified unique GO terms linked to ion transport and neurosecretory regulation. These pathways likely contribute to its local adaptation by maintaining ion homeostasis and enhancing the efficiency and reliability in neural signaling/synaptic transmission. The Aba population, meanwhile, possesses several unique GO terms related to muscle development and regulation of mating behavior in male bees (Figure 4C). This distinct adaptation strategy in Aba could further promote behavioral divergence, potentially leading to prezygotic isolation between Aba honeybees and other highland lineages (Arbuthnott, D., 2009; Matute, 2010; Ortiz‐Barrientos et al., 2009).

Notably, a large proportion of functional variants in each highland lineage are shared with polymorphisms present in lowland population (Figure 1C), which is consistent with the widespread mode of soft selection sweeps in the adaptive loci (Supplementary Table 2). This evolutionary mechanism supports the importance of standing genetic variation in rapid adaptation (Pool et al., 2017; Wallberg et al., 2017) as well as parallel evolution of insects (Chaturvedi et al., 2022; Lai et al., 2025; Montejo-Kovacevich et al., 2022). The critical role of standing genetic variation further imply that a population’s adaptive potential may be predictable based on its existing genetic repertoire.

### 4.2 Implications for adaptive prediction of populations to novel environments and conservation of population genomic variation

Maintaining the genetic diversity within and between populations of native, wild and domesticated species is a cornerstone for preserving their adaptive potential. This principle is central to both 2050 goals and 2030 targets in the Kunming-Montreal Global Biodiversity Framework (Convention on Biological Diversity, 2022). Considerable efforts have been devoted to modeling the evolutionary potential of natural populations to novel environmental stresses, including climate change. However, the accuracy of such models depends on the repeatability of evolutionary responses (e.g., deMayo and Ragland, 2025; Schlötterer, 2023). For complex polygenic traits like thermal adaptation, current predictive power is inherently limited due to widespread genetic redundancy in natural populations (Barghi et al., 2019; Burny et al., 2021; deMayo and Ragland, 2025). Research on parallel evolution in laboratory *Drosophila* populations further indicates that predicting the adaptive outcomes is challenging and requires a detailed knowledge of the adaptive architecture (with linkage disequilibrium) within the investigated populations (Schlötterer, 2023). However, the full adaptive architecture of a population could not be fully captured by any single study (Barghi et al., 2019), a limitation also evidenced by our findings of non-parallelism at adaptive loci among highland honeybee populations.

Moreover, conservation genetics in natural system should not only focus on the population or species itself, but also their interaction within the ecological network, which is important for biodiversity conservation in a functional ecosystem (Breed et al., 2019). Shift of adaptive loci toward the optimal fitness peak under novel stress will impact not only the target population/species, but would also indirectly impact biotic interactions in the local ecosystem. Recently, deMayo and colleagues (2025) highlighted the concerns that adaptation costs mediated by ecological processes in natural systems must be considered when predicting the adaptive potential. These adaptation costs are eco-evolutionary processes that can limit the extent of evolutionary rescue and reduce the adapted population’s fitness in alternative environments (Agrawal et al., 2010; deMayo et al., 2025; Hereford et al., 2009). A widely investigated system that demonstrating the cost of “adaptive” loci in local environment is that of the genetically modified (GM) crops in agriculture (e.g., Shelton et al., 2002). Research has showed that GM crops with different traits would exert different impacts on local ecosystem (Noack et al., 2024), implying the fitness of novel adaptation within a population/species may also be modulated by interactions with the local biota. Given the complexity of ecological networks, our efforts to conserve genetic variation should therefore account for both abiotic and biotic interactions. In turn, this integrated perspective will enhance the accuracy of predicting the adaptive potential of populations and species.

## Acknowledgments

We would like to thank the Information Technology Services in HKU and in NUS for offering research computing facilities in the computations of this project. We acknowledge the assistance of DeepSeek (https://chat.deepseek.com/) in writing scripts (in R or shell) used during the analysis of this research. The phenotype coding scheme was developed through discussions with DeepSeek. DeepSeek also assisted in generating the phenotype coding workflow and contributed to the English revision in part of the manuscript.

## Data and code availability

All sequencing data could be retrieved from public database and in our previous publication (Ke and Vasseur, 2026). Software and packages for analysis in this project are all available online. Shell and R scripts were attached in supplementary files.

## Author contributions

Conceptualization, F.S.K.; methodology and analysis, F.S.K.; Writing, F.S.K.

---

**Supplementary Figure 1.** Phenotypic convergence through directional parallel evolution. The pigmentation value scored on a scale of 0∼9 from black to light (Zhang et al., 2025).

**Supplementary Figure 2.** Super genetic network of 13 GO terms related to post-embryonic appendage and reproductive structure development via imaginal discs (Cluster A2 in Figure 2B). The pie chart indicates the presence of gene in each highland population.

**Supplementary Figure 3.** Degree distribution of the super genetic network (Supplementary Figure 2). Degree is number of direct connections that a gene has with other genes in the network.

**Supplementary Figure 4.** Jaccard Index of the adaptive genes in Cluster A2 of each population show a random distribution compared with the null distribution generated by 10,000 Monte Carlo simulations. Rikaze population shows a distinct pattern (e.g., Rikaze vs Aba, Rikaze vs Qinghai) likely support these adaptive genes may also interact with other pathways that further mediated local adaptation.

**Supplementary Figure 5.** Observed Jaccard Index of enriched GO terms and phenotypic traits compared with simulation. The null distribution was generated by 10,000 Monte Carlo simulations through randomly sampling the combined data with a constant size of each original set. The comparison between observed JI index and simulation across population pairs differentiated at both GO term and phenotypic levels, which is different from the pattern at the genetic level.

**Supplementary Table 1.** Sample information and bioclimate data of highland populations

**Supplementary Table 2.** Pipeline for adaptive loci filtering in each highland population

**Supplementary Table 3.** All identified genes potentially underlying highland adaptation of *Apis cerana*

**Supplementary Table 4.** Significantly enriched Gene Ontology (GO) terms of adaptive genes shared by two or more *A. cerana* highland populations (with *Drosophila melanogaster* database in GO annotation)

**Supplementary Table 5.** Significantly enriched GO terms of all adaptive genes (with *D. melanogaster* database)

**Supplementary Table 6.** Significantly enriched GO terms for each A. cerana highland population (with *D. melanogaster* database)

**Supplementary Table 7.** Morphological data of four highland and one lowland populations. Original data from Zhang et al., 2025.

**Supplementary Table 8.** Significantly enriched GO terms for each *A. cerana* highland population (with *A. cerana* database)

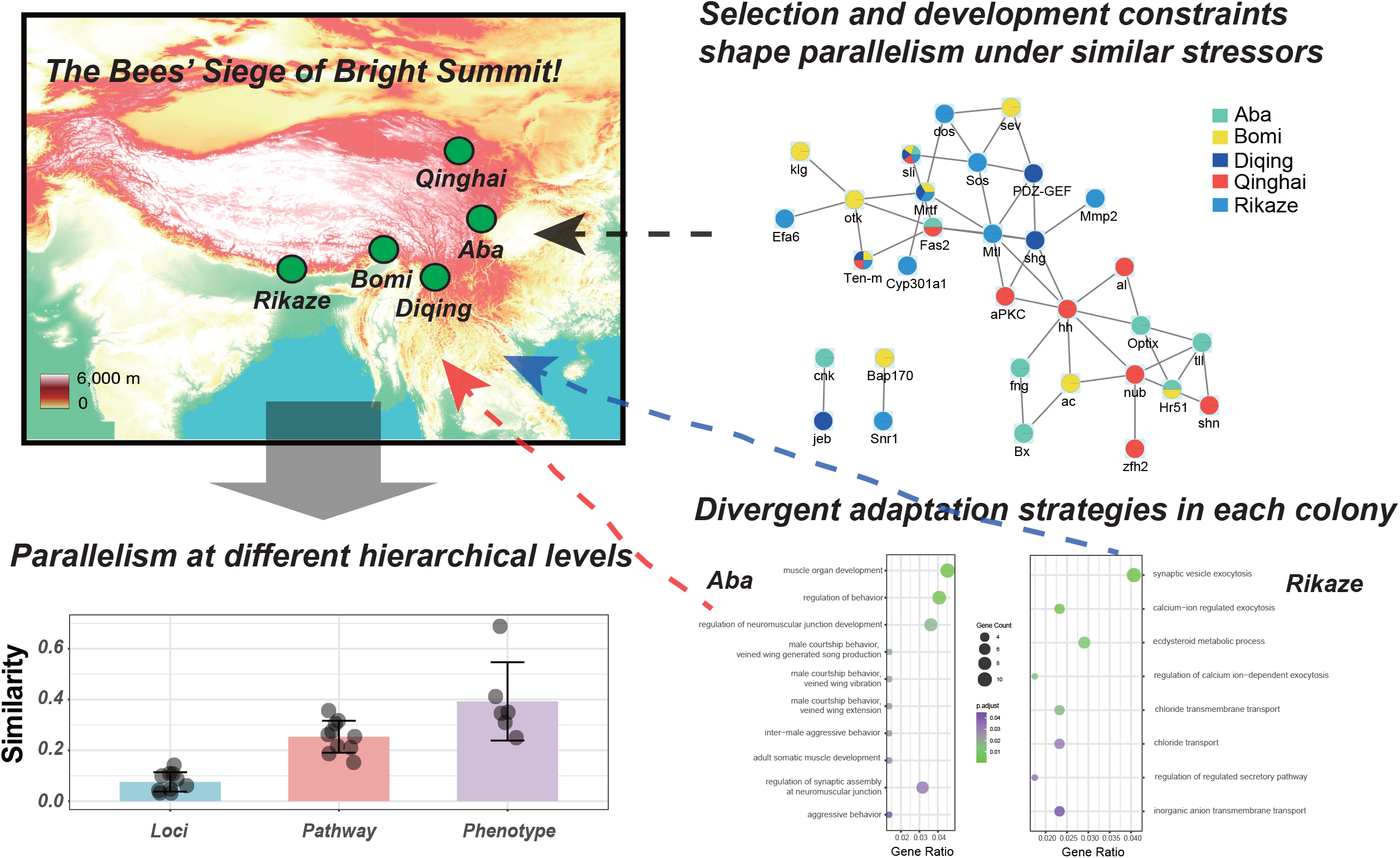

## Reference

Agrawal AA, Conner JK, Rasmann S. 2010. Tradeoffs and negative correlations in evolutionary ecology. In Evolution After Darwin: The First 150 Years, 243–268, Sinauer Associates.

Allard JB, Sharma S, Patel R, Sanderford M, Tamura K, Vucetic S, Gerhard GS, Kumar S. 2025. Evolutionary sparse learning reveals the shared genetic basis of convergent traits. Nat. Commun., 16(1): 3217.

Arbuthnott D. 2009. The genetic architecture of insect courtship behavior and premating isolation. Heredity, 103(1): 15–22.

Arnold SJ. 1992. Constraints on phenotypic evolution. Am. Nat., Suppl. 1: S85–107.

Babakhanlou M. 2018. Candidate gene identified in humans adapted to high-altitude depicts hypoxia tolerance in Drosophila melanogaster. PhD thesis, University of California, San Diego.

Barghi N, Schlötterer C. 2020. Distinct patterns of selective sweep and polygenic adaptation in evolve and resequence studies. Genome Biol. Evol., 12(6): 890–904.

Barghi N, Tobler R, Nolte V, Jakšić AM, Mallard F, Otte KA, Dolezal M, Taus T, Kofler R, Schlötterer C. 2019. Genetic redundancy fuels polygenic adaptation in Drosophila. PLoS Biol., 17(2): e3000128.

Barrett RD, Schluter D. 2008. Adaptation from standing genetic variation. Trends Ecol. Evol., 23(1): 38–44.

Bolnick DI, Barrett RD, Oke KB, Rennison DJ, Stuart YE. 2018. (Non) parallel evolution. Annu. Rev. Ecol. Evol. Syst., 49(1): 303–330.

Boyle EA, Li YI, Pritchard JK. 2017. An expanded view of complex traits: from polygenic to omnigenic. Cell, 169(7): 1177–1186.

Breed MF, Harrison PA, Blyth C, Byrne M, Gaget V, Gellie NJC, Groom SVC, Hodgson R, Mills JG, Prowse TAA, Steane DA, Mohr JJ. 2019. The potential of genomics for restoring ecosystems and biodiversity. Nat. Rev. Genet., 20(10): 615–628.

Brose K, Bland KS, Wang KH, Arnott D, Henzel W, Goodman CS, Tessier-Lavigne M, Kidd T. 1999. Slit proteins bind Robo receptors and have an evolutionarily conserved role in repulsive axon guidance. Cell, 96(6): 795–806.

Browning BL, Tian X, Zhou Y, Browning SR. 2021. Fast two-stage phasing of large-scale sequence data. Am. J. Hum. Genet., 108: 1880–1890.

Burny C, Nolte V, Dolezal M, Schlötterer C. 2021. Highly parallel genomic selection response in replicated Drosophila melanogaster populations with reduced genetic variation. Genome Biol. Evol., 13(11): evab239.

Cerca J. 2023. Understanding natural selection and similarity: Convergent, parallel and repeated evolution. Mol. Ecol., 32(20): 5451–5462.

Chang L, Zhu W, Chen Q, Zhao C, Sui L, Shen C, Zhang Q, Wang B, Jiang J. 2025. Adaptive divergence and functional convergence: The evolution of pulmonary gene expression in Amphibians of the Qingzang Plateau. Mol. Ecol., 34(5): e17663.

Chaturvedi S, Gompert Z, Feder JL, Osborne OG, Muschick M, Riesch R, Soria-Carrasco V, Nosil P. 2022. Climatic similarity and genomic background shape the extent of parallel adaptation in Timema stick insects. Nat. Ecol. Evol., 6(12): 1952–1964.

Cingolani P, Platts A, Wang le L, Coon M, Nguyen T, Wang L, Land SJ, Lu X, Ruden DM. 2012. A program for annotating and predicting the effects of single nucleotide polymorphisms, SnpEff: SNPs in the genome of Drosophila melanogaster strain w1118; iso-2; iso-3. Fly (Austin), 6(2): 80–92

Colosimo PF, Hosemann KE, Balabhadra S, Villarreal G Jr, Dickson M, Grimwood J, Schmutz J, Myers RM, Schluter D, Kingsley DM. 2005. Widespread parallel evolution in sticklebacks by repeated fixation of Ectodysplasin alleles. Science, 307(5717): 1928–1933.

Convention on Biological Diversity. 2022. Kunming-Montreal Global Biodiversity Framework. https://www.cbd.int/doc/c/e6d3/cd1d/daf663719a03902a9b116c34/cop-15-l-25-en.pdf.

Conte GL, Arnegard ME, Peichel CL, Schluter D. 2012. The probability of genetic parallelism and convergence in natural populations. Proc. Biol. Sci., 279(1749): 5039–5047.

Cresko WA, Amores A, Wilson C, Murphy J, Currey M, Phillips P, Bell MA, Kimmel CB, Postlethwait JH. 2004. Parallel genetic basis for repeated evolution of armor loss in Alaskan threespine stickleback populations. Proc. Natl. Acad. Sci., 101(16): 6050–6055.

Csilléry K, Rodríguez-Verdugo A, Rellstab C, Guillaume F. 2018. Detecting the genomic signal of polygenic adaptation and the role of epistasis in evolution. Mol. Ecol., 27(3): 606–612.

Danecek P, Auton A, Abecasis G, Albers CA, Banks E, DePristo MA, Handsaker RE, Lunter G, Marth GT, Sherry ST, McVean G, Durbin R; 1000 Genomes Project Analysis Group. 2011. The variant call format and VCFtools. Bioinformatics, 27(15): 2156–2158.

da Silva Ribeiro T, Lollar MJ, Sprengelmeyer QD, Huang Y, Benson DM, Orr MS, Johnson ZC, Corbett-Detig RB, Pool JE. 2025. Recombinant inbred line panels inform the genetic architecture and interactions of adaptive traits in Drosophila melanogaster. G3: Genes, Genom. Genet., 15(5): jkaf051.

DeGiorgio M, Huber CD, Hubisz MJ, Hellmann I, Nielsen R. 2016. SweepFinder2: increased sensitivity, robustness and flexibility. Bioinformatics, 32(12): 1895–1897.

deMayo JA, Ragland GJ. 2025. (Limited) Predictability of thermal adaptation in invertebrates. J. Exp. Biol., 228(5): JEB249450.

de Vladar HP, Barton N. 2014. Stability and response of polygenic traits to stabilizing selection and mutation. Genetics, 197(2): 749–767.

Ding D, Liu G, Hou L, Gui W, Chen B, Kang L. 2018. Genetic variation in PTPN1 contributes to metabolic adaptation to high-altitude hypoxia in Tibetan migratory locusts. Nat. Commun., 9(1): 4991.

Ellers J, Boggs CL. 2004. Functional ecological implications of intraspecific differences in wing melanization in Colias butterflies. Biol. J. Linn. Soc., 82(1): 79–87.

Feng S, DeGrey SP, Guédot C, Schoville SD, Pool JE. 2024. Genomic diversity illuminates the environmental adaptation of Drosophila suzukii. Genome Biol. Evol., 16(9): evae195.

Garud NR, Messer P, Buzbas E, and Petrov D. 2015. Soft selective sweeps are the primary mode of adaptation in Drosophila. PLoS Genet., 11: e1005004.

Hallgrimsson B, Green RM, Katz DC, Fish JL, Bernier FP, Roseman CC, Young NM, Cheverud JM, Marcucio RS. 2019. The developmental-genetics of canalization. Semin. Cell Dev. Biol., 88: 67–79.

Hämälä T, Gorton AJ, Moeller DA, Tiffin P. 2020. Pleiotropy facilitates local adaptation to distant optima in common ragweed (Ambrosia artemisiifolia). PLoS genet., 16(3): e1008707.

Hereford J. 2009. A quantitative survey of local adaptation and fitness trade-offs. Am. Nat., 173: 579–588

Höllinger I, Pennings PS, Hermisson J. 2019. Polygenic adaptation: from sweeps to subtle frequency shifts. PLoS Genet., 15(3): e1008035.

Huerta-Cepas J, Forslund K, Coelho LP, Szklarczyk D, Jensen LJ, von Mering C, Bork P. 2017. Fast genome-wide functional annotation through orthology assignment by eggNOG-mapper. Mol. Biol. Evol., 34(8): 2115–2122.

Huerta-Cepas J, Szklarczyk D, Heller D, Hernández-Plaza A, Forslund SK, Cook H, Mende DR, Letunic I, Rattei T, Jensen LJ, von Mering C, Bork P. 2019. eggNOG 5.0: a hierarchical, functionally and phylogenetically annotated orthology resource based on 5090 organisms and 2502 viruses. Nucleic Acids Res., 47(D1): D309–D314.

Ji Y, Li X, Ji T, Tang J, Qiu L, Hu J, Dong J, Luo S, Liu S, Frandsen PB, Zhou X. 2020. Gene reuse facilitates rapid radiation and independent adaptation to diverse habitats in the Asian honeybee. Sci. Adv., 6(51): eabd3590.

Ke F, Vasseur L. 2026. Contribution of local recombination and AT-biased mutations to differentiated region formation in Apis cerana.

Lai WY, Hsu SK, Futschik A, Schlötterer C., 2025. Pleiotropy increases parallel selection signatures during adaptation from standing genetic variation. eLife, 13: RP102321.

Lipshutz SE, Hibbins MS, Bentz AB, Buechlein AM, Empson TA, George EM, Hauber ME, Rusch DB, Schelsky WM, Thomas QK, Torneo SJ, Turner AM, Wolf SE, Woodruff MJ, Hahn MW, Rosvall KA. 2025. Repeated behavioural evolution is associated with convergence of gene expression in cavity-nesting songbirds. Nat. Ecol. Evol., 9: 845–856.

Losos JB. 2011. Convergence, adaptation, and constraint. Evolution, 65(7): 1827–40.

Mani MS. 1968. Ecological specializations of high altitude insects. In Ecology and biogeography of high altitude insects. Series Entomologica, Volume 4: 51–74. Springer Netherlands.

Matute DR, 2010. Reinforcement of gametic isolation in Drosophila. PLoS Biol., 8(3): e1000341.

Montero-Mendieta S, Tan K, Christmas MJ, Olsson A, Vilà C, Wallberg A, Webster MT. 2019. The genomic basis of adaptation to high‐altitude habitats in the eastern honey bee (Apis cerana). Mol. Ecol., 28(4): 746–760.

Montejo-Kovacevich G, Meier JI, Bacquet CN, Warren IA, Chan YF, Kucka M, Salazar C, Rueda-M N, Montgomery SH, McMillan WO, Kozak KM, Nadeau NJ, Martin SH, Jiggins CD. 2022. Repeated genetic adaptation to altitude in two tropical butterflies. Nat. Commun., 13(1): 4676.

Noack F, Engist D, Gantois J, Gaur V, Hyjazie BF, Larsen A, M’Gonigle LK, Missirian A, Qaim M, Sargent RD, Souza-Rodrigues E, Kremen C. 2024. Environmental impacts of genetically modified crops. Science, 385(6712): eado9340.

Ortiz-Barrientos D, Grealy A, Nosil P. 2009. The genetics and ecology of reinforcement: implications for the evolution of prezygotic isolation in sympatry and beyond. Ann. N. Y. Acad Sci., 1168(1): 156–182.

Pamenter ME, Hall JE, Tanabe Y, Simonson TS. 2020. Cross-species insights into genomic adaptations to hypoxia. Front. Genet., 11: 743.

Parkash R, Rajpurohit S, Ramniwas S. 2008. Changes in body melanisation and desiccation resistance in highland vs. lowland populations of D. melanogaster. J. Insect Physiol., 54(6): 1050–1056.

Pfenninger M, Patel S, Arias-Rodriguez L, Feldmeyer B, Riesch R, Plath M. 2015. Unique evolutionary trajectories in repeated adaptation to hydrogen sulphide-toxic habitats of a neotropical fish (Poecilia mexicana). Mol. Ecol., 24(21): 5446–5459.

Pickersgill B. 2018. Parallel vs. convergent evolution in domestication and diversification of crops in the Americas. Front. Ecol. Evol., 6: 56.

Pool JE, Braun DT, Lack JB. 2017. Parallel evolution of cold tolerance within Drosophila melanogaster. Mol Biol Evol., 34(2): 349–360.

Reed RD, Papa R, Martin A, Hines HM, Counterman BA, Pardo-Diaz C, Jiggins CD, Chamberlain NL, Kronforst MR, Chen R, Halder G, Nijhout HF, McMillan WO. 2011. optix drives the repeated convergent evolution of butterfly wing pattern mimicry. Science, 333(6046): 1137–1141.

Rosenblum EB, Parent CE, and Brandt EE. 2014. The molecular basis of phenotypic convergence. Annu. Rev. Ecol. Evol. S., 45: 203–226.

Rothberg JM, Jacobs JR, Goodman CS, Artavanis-Tsakonas S. 1990. slit: an extracellular protein necessary for development of midline glia and commissural axon pathways contains both EGF and LRR domains. Genes Dev., 4(12A): 2169–2187.

R Core Team. 2023. R: A Language and Environment for Statistical Computing. R Foundation for Statistical Computing, Vienna, Austria.

Sackton TB, Clark N. 2019. Convergent evolution in the genomics era: new insights and directions. Philos. Trans. R. Soc. Lond. B. Biol. Sci., 374(1777): 20190102.

Schlötterer C. 2023. How predictable is adaptation from standing genetic variation? Experimental evolution in Drosophila highlights the central role of redundancy and linkage disequilibrium. Phil. Trans. R. Soc. B., 378 (1877): 20220046.

Schwenk K, Wagner GP. 2001. Function and the evolution of phenotypic stability: connecting pattern to process. Am. Zool., 41(3): 552–563.

Shakya SB, Edwards SV, Sackton TB. 2025. Convergent evolution of noncoding elements associated with short tarsus length in birds. BMC Biol. 23(1): 52.

Shannon P, Markiel A, Ozier O, Baliga NS, Wang JT, Ramage D, Amin N, Schwikowski B, Ideker T. 2003. Cytoscape: a software environment for integrated models of biomolecular interaction networks. Genome Res., 13(11): 2498–2504.

Sharma V, Varshney R, Sethy NK. 2022. Human adaptation to high altitude: a review of convergence between genomic and proteomic signatures. Hum. Genomics, 16: 21.

Shelton AM, Zhao JZ, Roush RT. 2002. Economic, ecological, food safety, and social consequences of the deployment of bt transgenic plants. Annu. Rev. Entomol., 47: 845–881.

Shi YY, Sun LX, Huang ZY, Wu XB, Zhu YQ, Zheng HJ, Zeng ZJ. 2013. A SNP based high-density linkage map of Apis cerana reveals a high recombination rate similar to Apis mellifera. PLoS One, 8(10): e76459.

Sow MD, Forestan C, Pont C, Civan P, Battaglia R, Seidel M, Siguret C, Curci PL, Tondelli A, Bustos Korts D, Mazzucotelli E, Leroy T, Huneau C, Delahaye M, Ormanbekova D, Bozzoli M, Guarino-Vignon P, Schaal C, Cabanis M, Lelievre M, Cayrol J, Guerra D, Nigro D, Gadaleta A, Ens J, Wiebe K, Shapiro B, Green RE, van Eeuwijk F, Bayer M, Russell J, Dawson I, Waugh R, Kilian B, Orlando L, Sonnante G, Pozniak CJ, Tuberosa R, Haberer G, Maccaferri M, Cattivelli L, Salse J. 2025. Striking convergent selection history of wheat and barley and its potential for breeding. Nat. Plants, 11(11): 2268–2285.

Stern DL. 2013. The genetic causes of convergent evolution. Nat. Rev. Genet., 14(11): 751–764.

Storz JF, Cheviron ZA. 2021. Physiological genomics of adaptation to high-altitude hypoxia. Annu. Rev. Anim. Biosci., 9: 149–171.

Storz JF, Scott GR. 2019. Life ascending: mechanism and process in physiological adaptation to high-altitude hypoxia. Annu. Rev. Ecol. Evol. Syst., 50(1): 503–526.

Su YC, Chiu YF, Warrit N, Otis GW, Smith DR. 2023. Phylogeography and species delimitation of the Asian cavity-nesting honeybees. Insect Syst. Diversity, 7(4): 5.

Swaminathan A, Xia F, Rohner N. 2024. From darkness to discovery: evolutionary, adaptive, and translational genetic insights from cavefish. Trends Genet., 40(1): 24–38.

Thorhölludottir DAV, Hsu SK, Barghi N, Mallard F, Nolte V, Schlötterer C. 2025. Reduced parallel gene expression evolution with increasing genetic divergence-a hallmark of polygenic adaptation. Mol. Ecol., 34(12): e17803.

Tishkoff SA, Reed FA, Ranciaro A, Voight BF, Babbitt CC, Silverman JS, Powell K, Mortensen HM, Hirbo JB, Osman M, Ibrahim M, Omar SA, Lema G, Nyambo TB, Ghori J, Bumpstead S, Pritchard JK, Wray GA, Deloukas P. 2007. Convergent adaptation of human lactase persistence in Africa and Europe. Nat. Genet., 39(1): 31–40.

True JR, Haag ES. 2001. Developmental system drift and flexibility in evolutionary trajectories. Evol. Dev., 3(2): 109–119.

Wallberg A, Schöning C, Webster MT, Hasselmann M. 2017. Two extended haplotype blocks are associated with adaptation to high altitude habitats in East African honey bees. PLoS Genet., 13(5): e1006792.

Wang Y, McNeil P, Abdulazeez R, Pascual M, Johnston SE, Keightley PD, Obbard DJ. 2023. Variation in mutation, recombination, and transposition rates in Drosophila melanogaster and Drosophila simulans. Genome Res., 33(4): 587–598.

Wickham H, Averick M, Bryan J, Chang W, McGowan LD, François R, Grolemund G, Hayes A, Henry L, Hester J, Kuhn M, Pedersen TL, Miller E, Bache SM, Müller K, Ooms J, Robinson D, Seidel DP, Spinu V, Takahashi K, Vaughan D, Wilke C, Woo K, Yutani H. 2019. Welcome to the tidyverse. J. Open Source Softw., 4(43): 1686.

Wickham H. 2016. Ggplot2: Elegant graphics for data analysis. 2nd ed. Cham., Switzerland, Springer International Publishing.

Wong K, Park HT, Wu JY, Rao Y. 2002. Slit proteins: molecular guidance cues for cells ranging from neurons to leukocytes. Curr. Opin. Genet. Dev., 12(5): 583–591.

Wu DD, Yang CP, Wang MS, Dong KZ, Yan DW, Hao ZQ, Fan SQ, Chu SZ, Shen QS, Jiang LP, Li Y, Zeng L, Liu HQ, Xie HB, Ma YF, Kong XY, Yang SL, Dong XX, Esmailizadeh A, Irwin DM, Xiao X, Li M, Dong Y, Wang W, Shi P, Li HP, Ma YH, Gou X, Chen YB, Zhang YP. 2020. Convergent genomic signatures of high-altitude adaptation among domestic mammals. Natl. Sci. Rev., 7(6): 952–963.

Wu T, Hu E, Xu S, Chen M, Guo P, Dai Z, Feng T, Zhou L, Tang W, Zhan L, Fu X, Liu S, Bo X, Yu G. 2021. clusterProfiler 4.0: A universal enrichment tool for interpreting omics data. Innovation (Camb)., 2(3): 100141.

Wu W, Wong K, Chen J, Jiang Z, Dupuis S, Wu JY, Rao Y. 1999. Directional guidance of neuronal migration in the olfactory system by the protein Slit. Nature, 400(6742): 331–336.

Xu S, He Z, Guo Z, Zhang Z, Wyckoff GJ, Greenberg A, Wu CI, Shi S. 2017. Genome-wide convergence during evolution of mangroves from woody plants. Mol. Biol. Evol., 34(4): 1008–1015.

Yeaman S, Hodgins KA, Lotterhos KE, Suren H, Nadeau S, Degner JC, Nurkowski KA, Smets P, Wang T, Gray LK, Liepe KJ, Hamann A, Holliday JA, Whitlock MC, Rieseberg LH, Aitken SN. 2016. Convergent local adaptation to climate in distantly related conifers. Science, 353(6306): 1431–1433.

Yeaman S. 2022. Evolution of polygenic traits under global vs local adaptation. Genetics, 220(1): iyab134.

Zhang J, Zhang X, Liu N, Hu J, Hiller M, Sharma V, Han F, Dai H, Tu X, Cooper DN, Wu DD, Zeng L. 2026. A POMT2 missense substitution contributes to hypoxia adaptation in hibernating mammals. Mol. Biol. Evol., msag001.

Zhang Q, Gou W, Wang X, Zhang Y, Ma J, Zhang H, Zhang Y, Zhang H. 2016. Genome resequencing identifies unique adaptations of Tibetan chickens to hypoxia and high-dose ultraviolet radiation in high-altitude environments. Genome Biol. Evol., 8(3): 765–776.

Zhang T, Chen J, Zhang J, Guo YT, Zhou X, Li MW, Zheng ZZ, Zhang TZ, Murphy RW, Nevo E, Shi P. 2021. Phenotypic and genomic adaptations to the extremely high elevation in plateau zokor (Myospalax baileyi). Mol. Ecol., 30(22): 5765–5779.

Zhang X, Lu J, Qu X, Chen X. 2025. An evaluation of morphometric characteristics of honey bee (Apis cerana) populations in the Qinghai–Tibet Plateau in China. Life, 15: 255.

Zhou D, Azad P, Stobdan T, Haddad GG. 2021. Mechanisms regulating hypoxia tolerance in Drosophila and humans. In Stress: Genetics, Epigenetics and Genomics, Volume 4: 241–251. Academic Press.

