## Supplementary Figure 1-5 for "Understanding highland adaptation of *Apis cerana* through repeated while independent colonization"

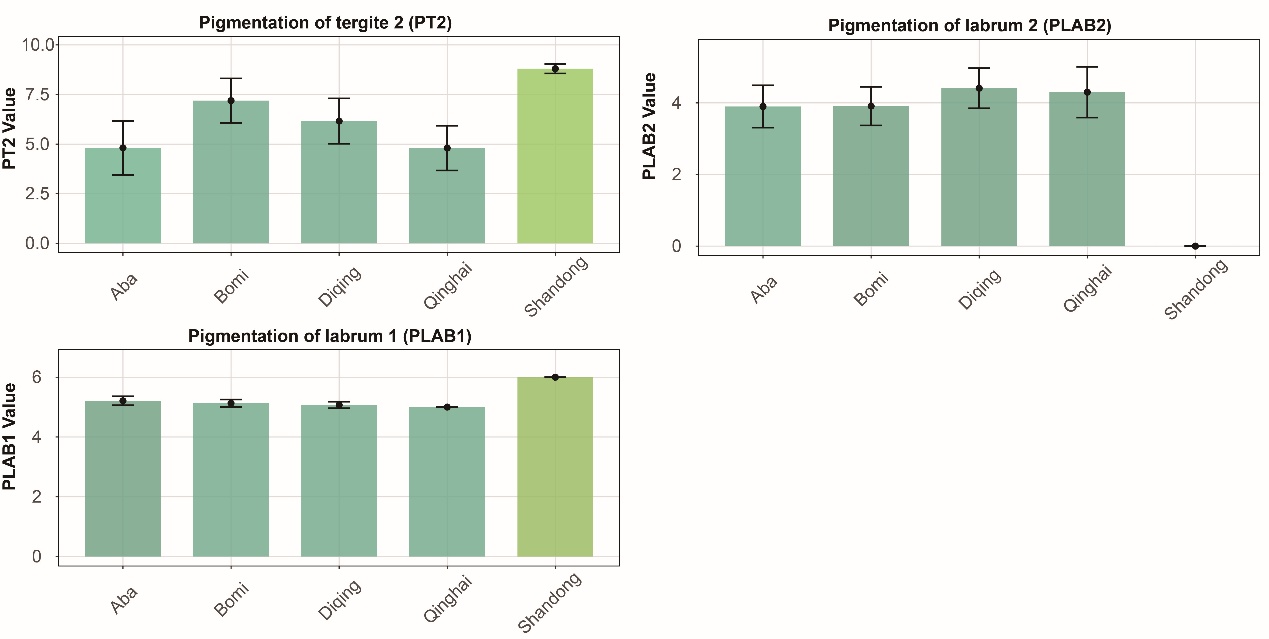

**Supplementary Figure 1.** Phenotypic convergence through directional parallel evolution. The pigmentation value scored on a scale of 0~9 from black to light (Zhang et al., 2025).

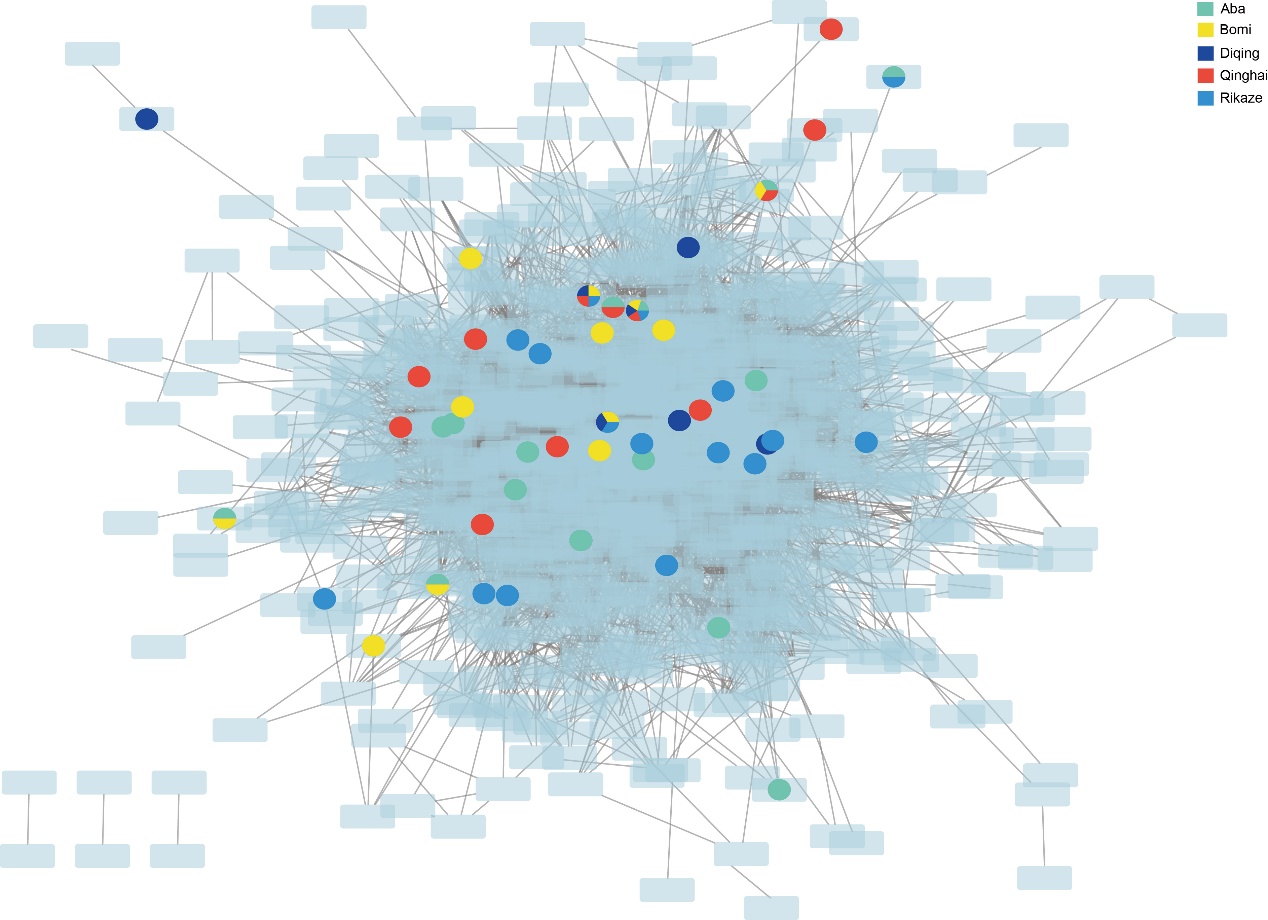

**Supplementary Figure 2**. Super genetic network of 13 GO terms related to post-embryonic appendage and reproductive structure development via imaginal discs (Cluster A2 in Figure 2B). The pie chart indicates the presence of gene in each highland population.

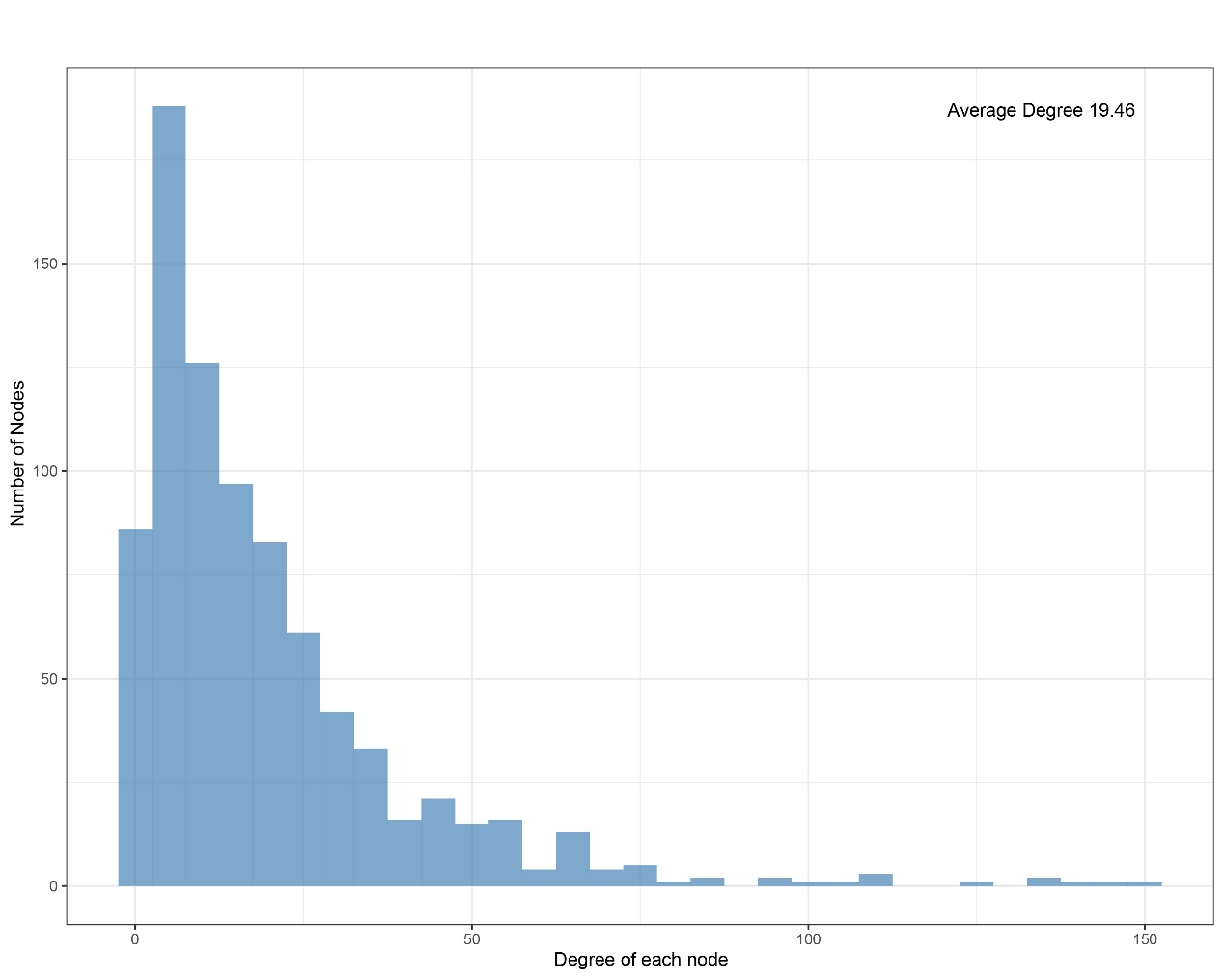

**Supplementary Figure 3**. Degree distribution of the super genetic network (Supplementary Figure 2). Degree is number of direct connections that a gene has with other genes in the network.

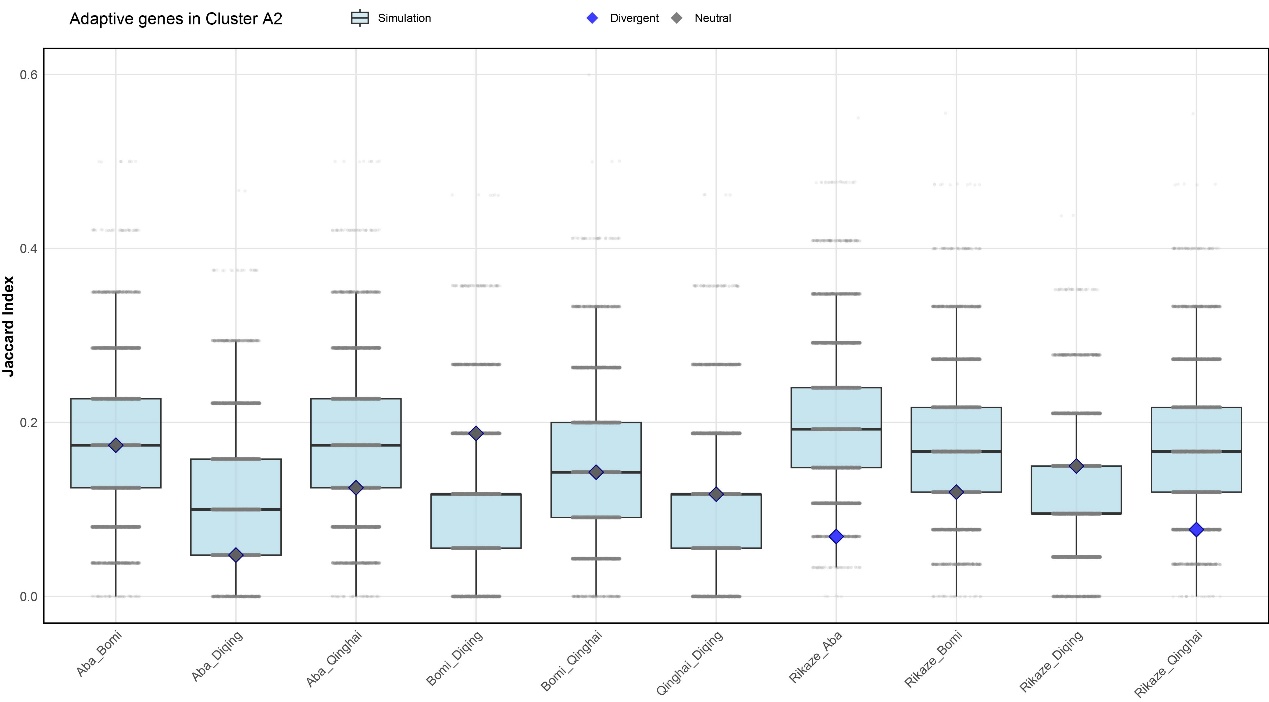

**Supplementary Figure 4**. Jaccard Index of the adaptive genes in Cluster A2 of each population show a random distribution compared with the null distribution generated by 10,000 Monte Carlo simulations. Rikaze population shows a distinct pattern (e.g., Rikaze vs Aba, Rikaze vs Qinghai) likely support these adaptive genes may also interact with other pathways that further mediated local adaptation.

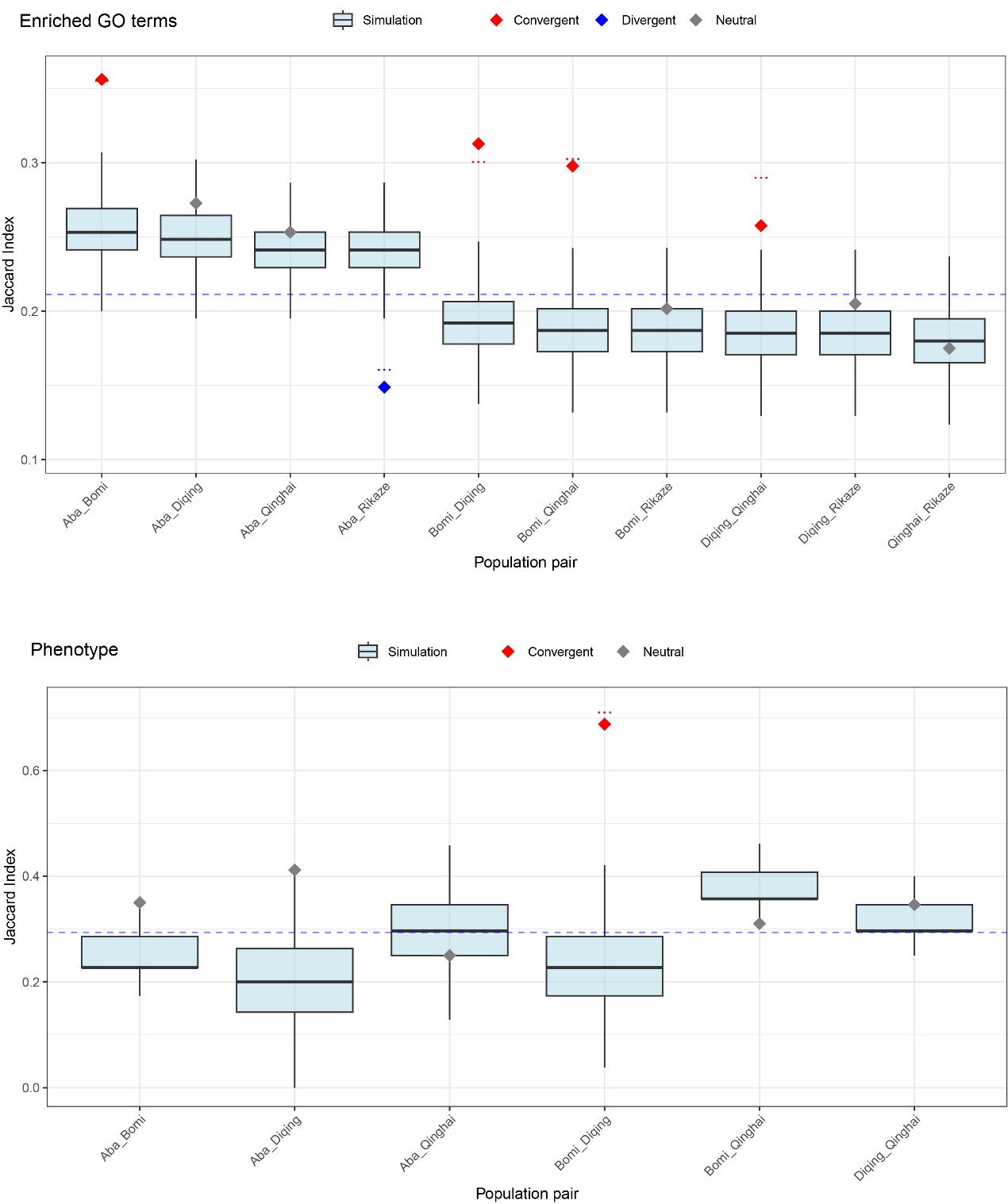

**Supplementary Figure 5**. Observed Jaccard Index of enriched GO terms and phenotypic traits compared with simulation. The null distribution was generated by 10,000 Monte Carlo simulations through randomly sampling the combined data with a constant size of each original set. The comparison between observed JI index and simulation across population pairs differentiated at both GO term and phenotypic levels, which is different from the pattern at the genetic level.
